## Supplementary Material for "Multi-level processing of emotions in life motion signals revealed through pupil responses"

**The motion stimuli used in Experiments 1-4.**

**Experiment 1: Intact BM**

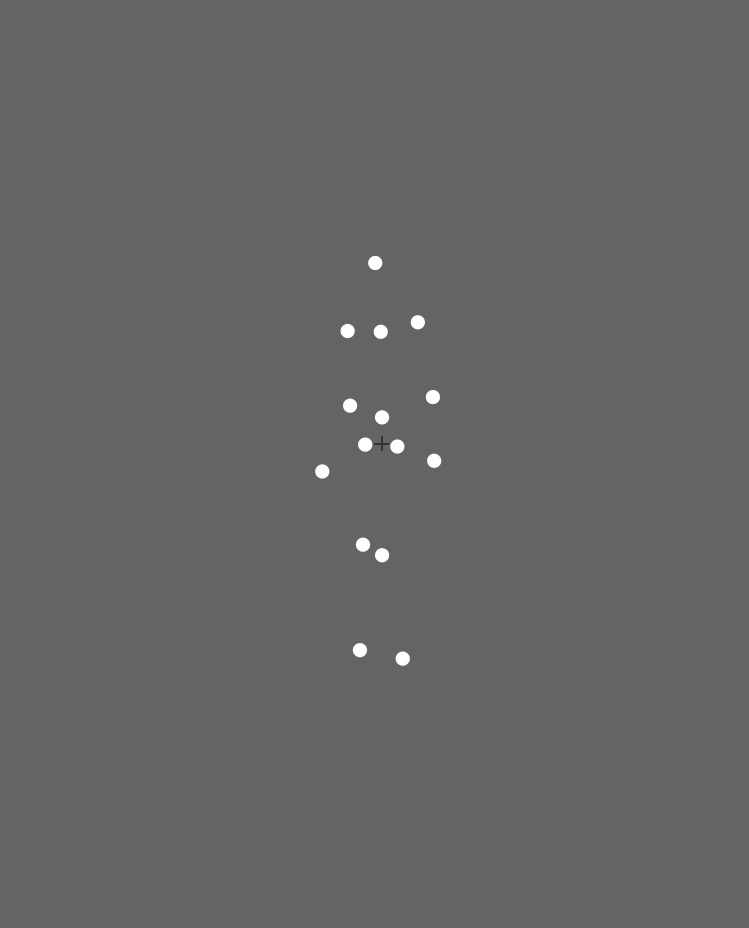

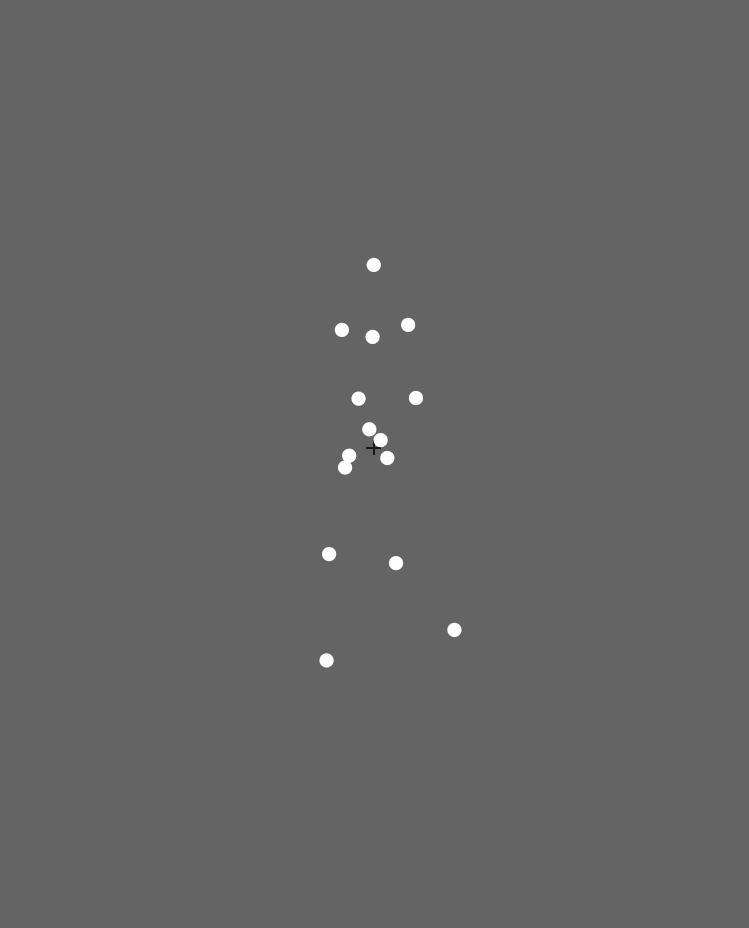

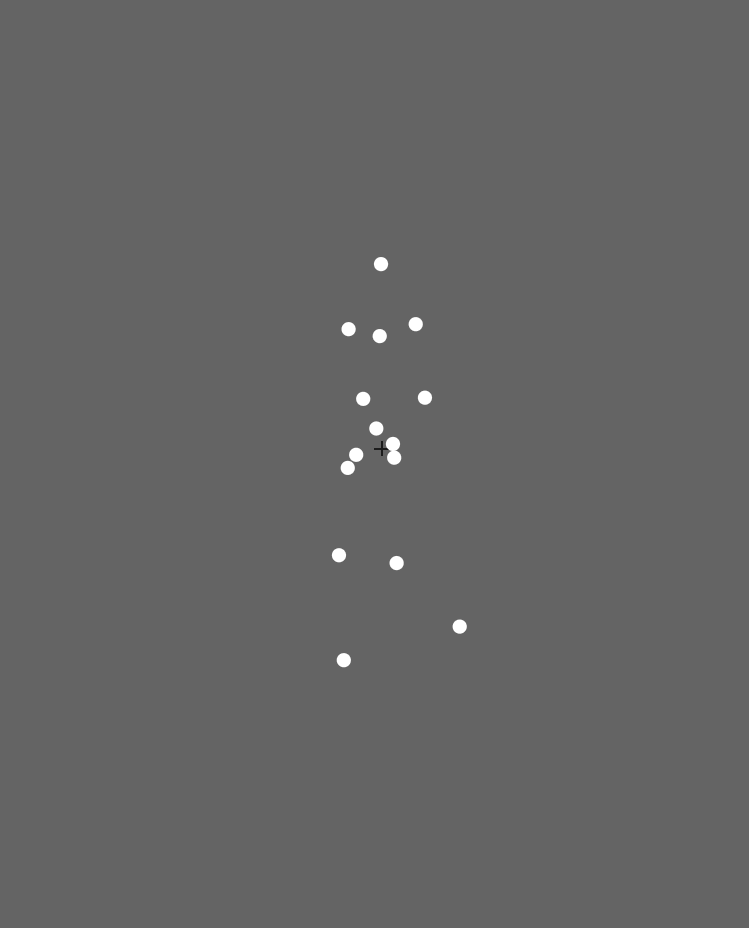

sad

neutral

happy

**Experiment 2: Inverted BM**

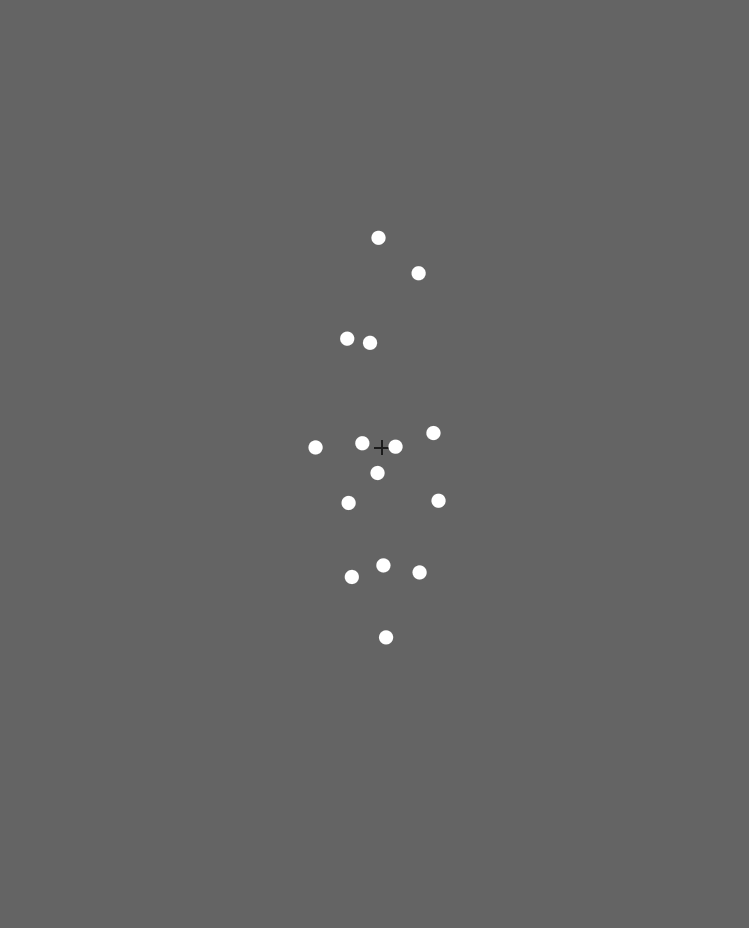

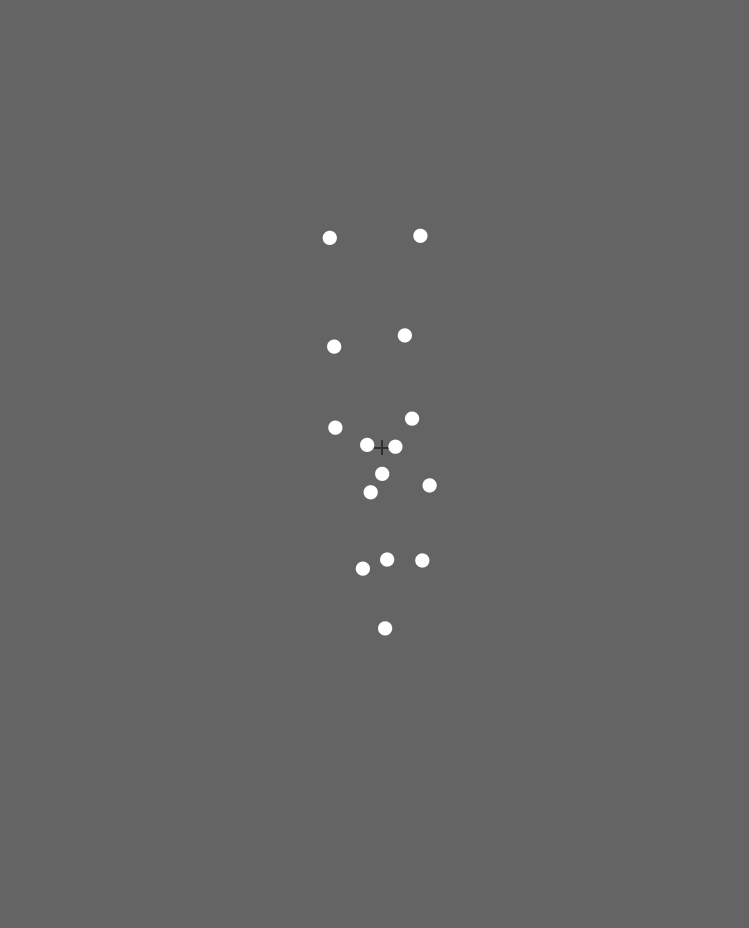

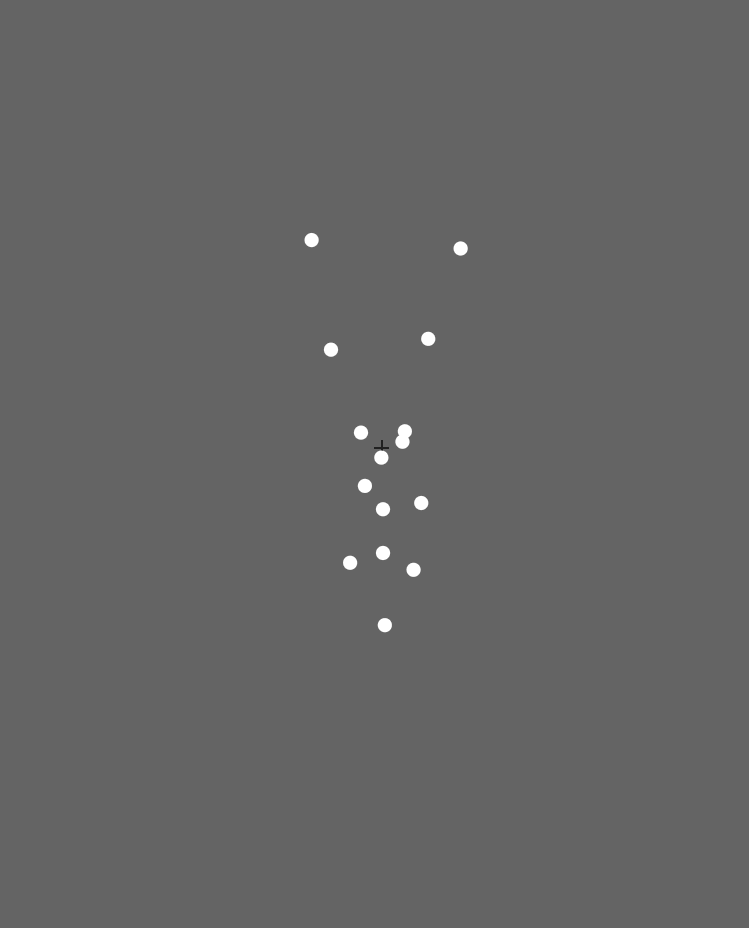

sad

neutral

happy

**Experiment 3: Nonbiological motion**

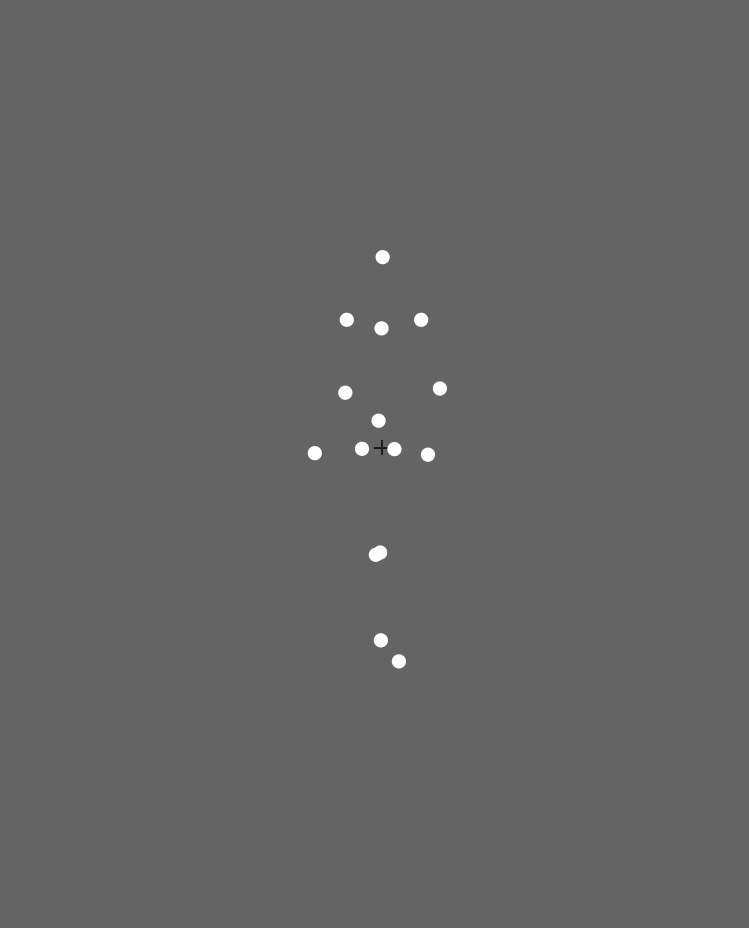

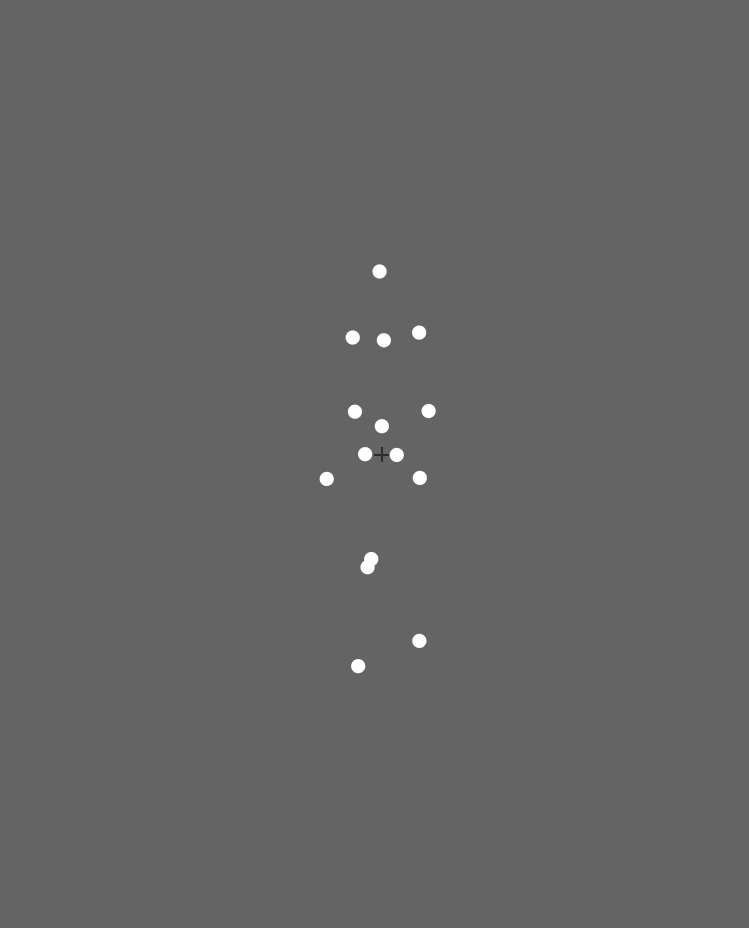

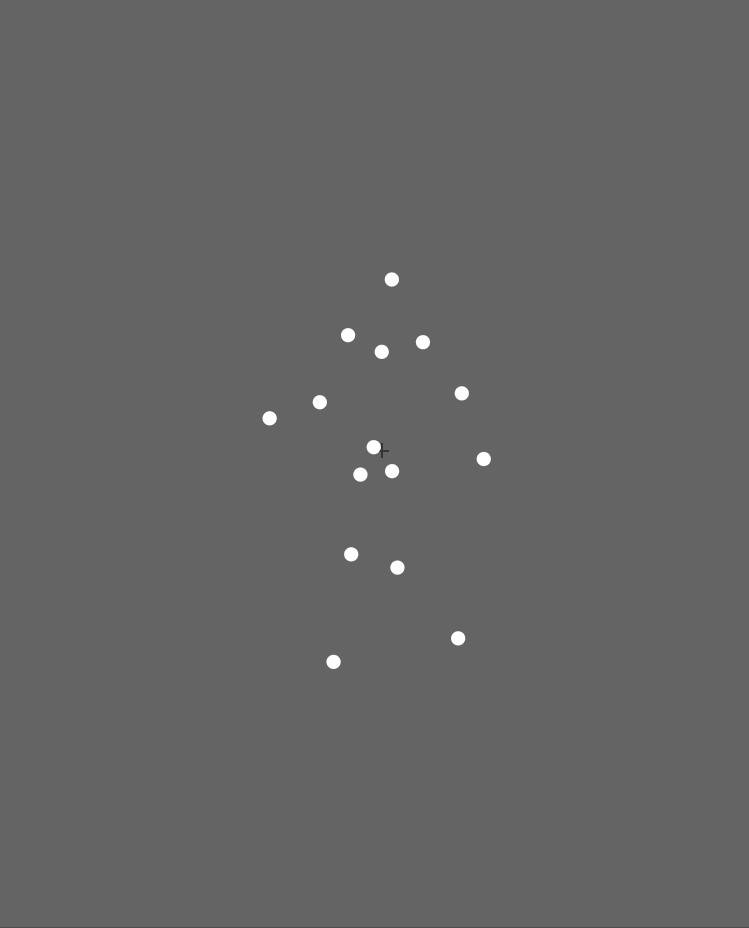

neutral

happy

sad

**Experiment 4: Local BM**

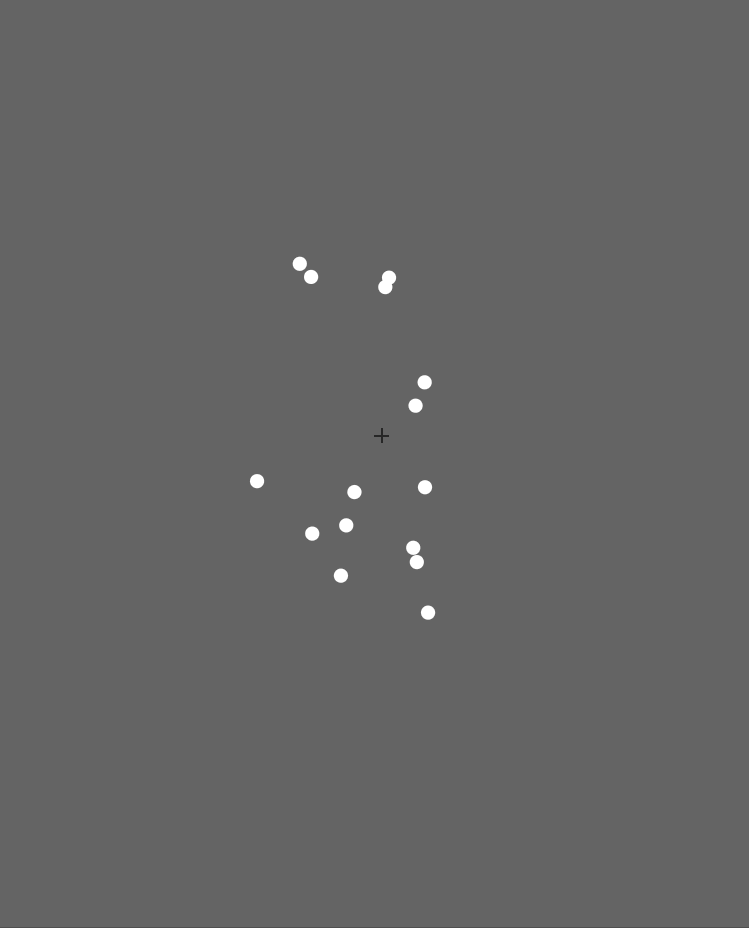

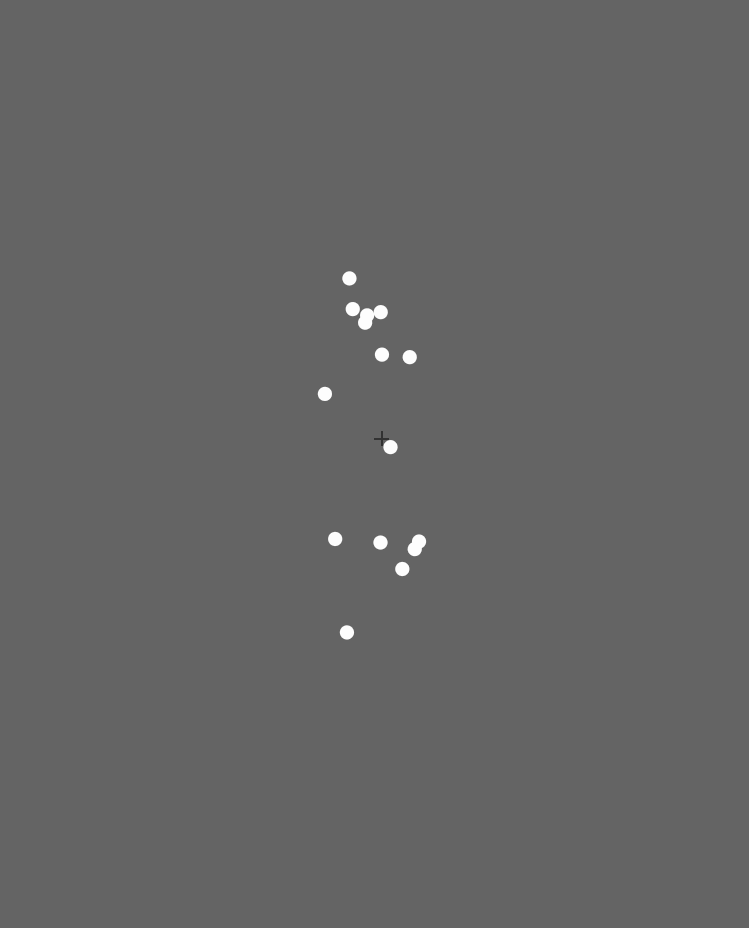

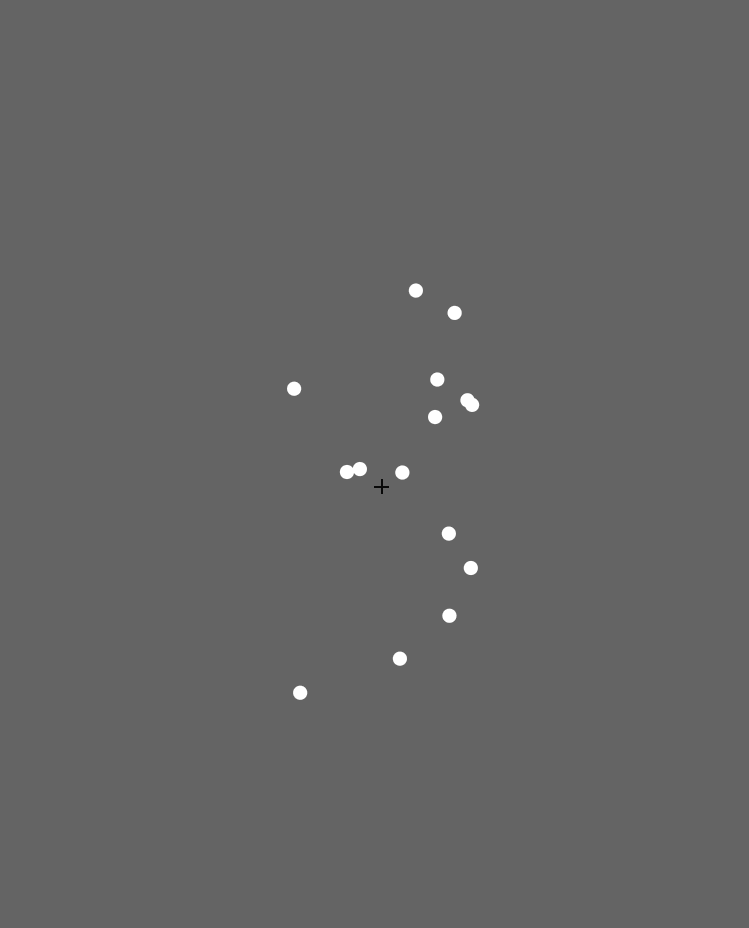

neutral

sad

happy

**Time-course results of Experiment 1b**

To strengthen the results of Experiment 1a, a replication experiment (Experiment 1b) was conducted and we added a retest to further assess the reliability of the results. The cluster-based permutation analysis of the first test of Experiment 1b revealed that the happy BM induced significantly larger pupil responses than the neutral BM from 3250 ms to 4000 ms, and the sad BM evoked significantly smaller pupil response than the neutral BM from 2950-4000 ms. Additionally, the happy BM evoked a significantly larger pupil response as compared to the sad BM from 1450 ms to 4000 ms (see Fig. 1A). Results of the second test revealed that the happy BM induced a significant pupil dilation effect than the neutral BM from 1900-4000 ms, and the sad BM evoked a significantly smaller pupil response than the neutral BM from 2450-4000 ms. Additionally, the happy BM evoked larger pupil responses as compared to the sad BM from from 1200 ms to 4000 ms (see Fig. 1B).

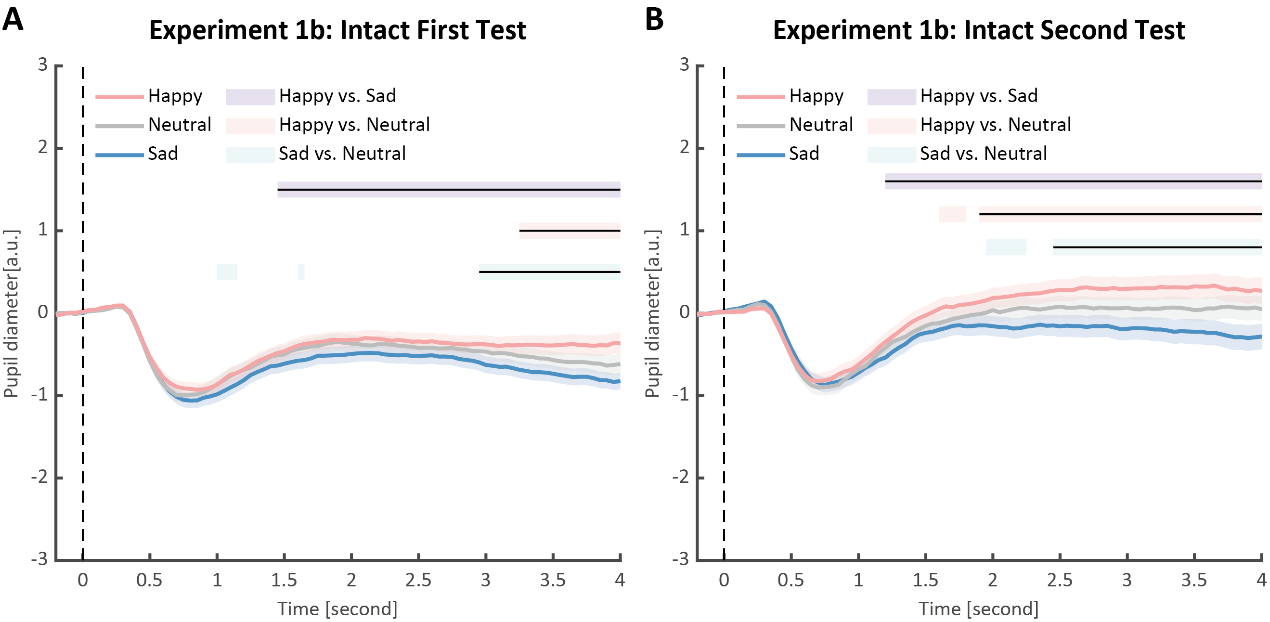

**Fig. 1.** Time course of pupil responses to happy, sad, and neutral BM in Experiment 1b. Solid lines represent pupil diameter under each emotional condition as a function of time (happy: red; sad: blue; neutral: gray); shaded areas represent the SEM between participants; colored horizontal lines indicate periods during which there are statistically significant differences among conditions at *p* <0.05; and black horizontal lines indicate significant differences after cluster-based permutation correction. All the pupil data are in arbitrary units (a.u.). (A) In the first test of Experiment 1b, we successfully replicated the results of Experiment 1a: the happy BM evoked larger pupil response as compared to the sad and neutral BM, and the sad BM evoked smaller pupil size than the neutral BM. (B) such results were similarly observed in the retest.

**Correlation results between AQ and the pupil responses induced by happy/sad/neutral BM**

No significant correlations were found between AQ and the original pupil responses across four experiments (see Fig.2). This is potentially because the original pupil response is a mixed result of stimuli perception and emotion perception, while the pupil changes across emotional conditions could more faithfully reflect individual sensitivities to emotions in BM (Burley et al., 2017; Pomè et al., 2020; Turi et al., 2018).

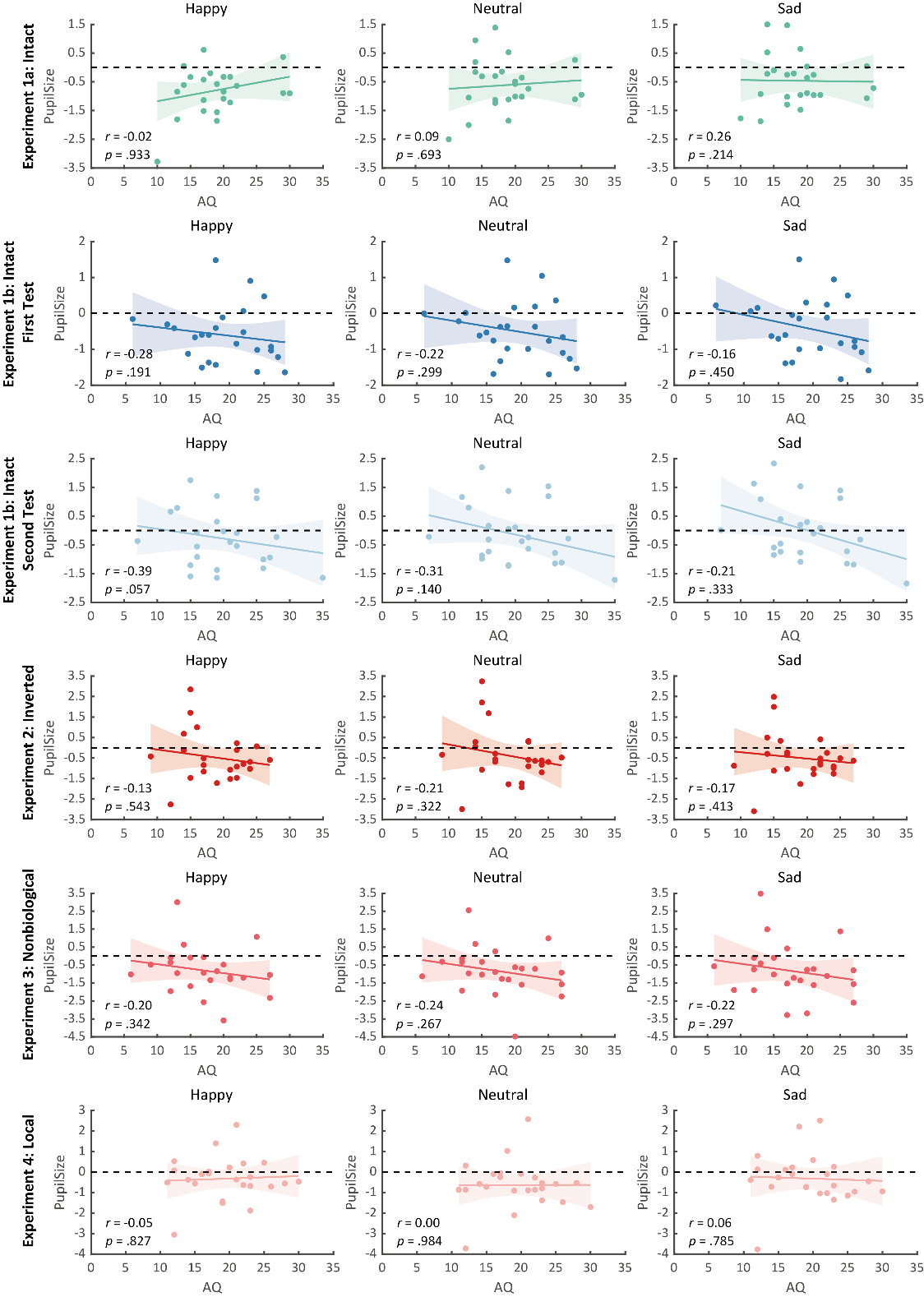

**Fig. 2.** Correlation results for AQ scores and pupil responses towards happy, sad, and neutral BM across four experiments. No significant correlations were observed. Dots in the scatter plot indicate the individual data and the shaded region indicate the 95% confidence interval.

**References:**

| Burley, D. T., Gray, N. S., & Snowden, R. J. (2017). As Far as the Eye Can See: Relationship between Psychopathic Traits and Pupil Response to Affective Stimuli. *PLOS ONE, 12*(1), e0167436. https://doi.org/10.1371/journal.pone.0167436  Pomè, A., Binda, P., Cicchini, G. M., & Burr, D. C. (2020). Pupillometry correlates of visual priming, and their dependency on autistic traits. *Journal of vision, 20*(3), 3-3. |
| --- |
| Turi, M., Burr, D. C., & Binda, P. (2018). Pupillometry reveals perceptual differences that are tightly linked to autistic traits in typical adults. *eLife*, *7*. https://doi.org/10.7554/elife.32399 |
