## Supplementary figures and images for "Multi-level processing of emotions in life motion signals revealed through pupil responses"

### Supplemenatry Figure 1

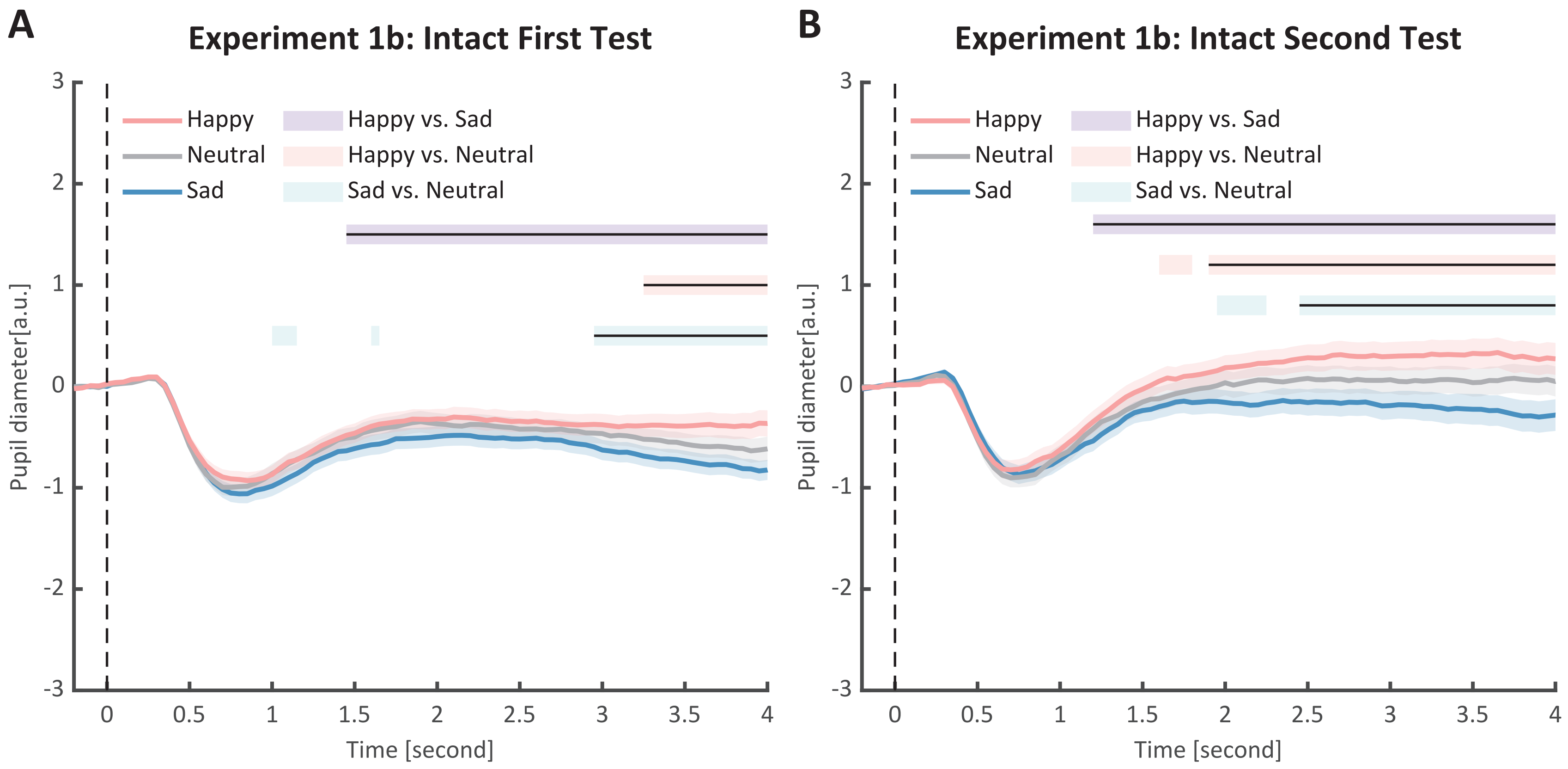

### Supplemenatry Figure 2

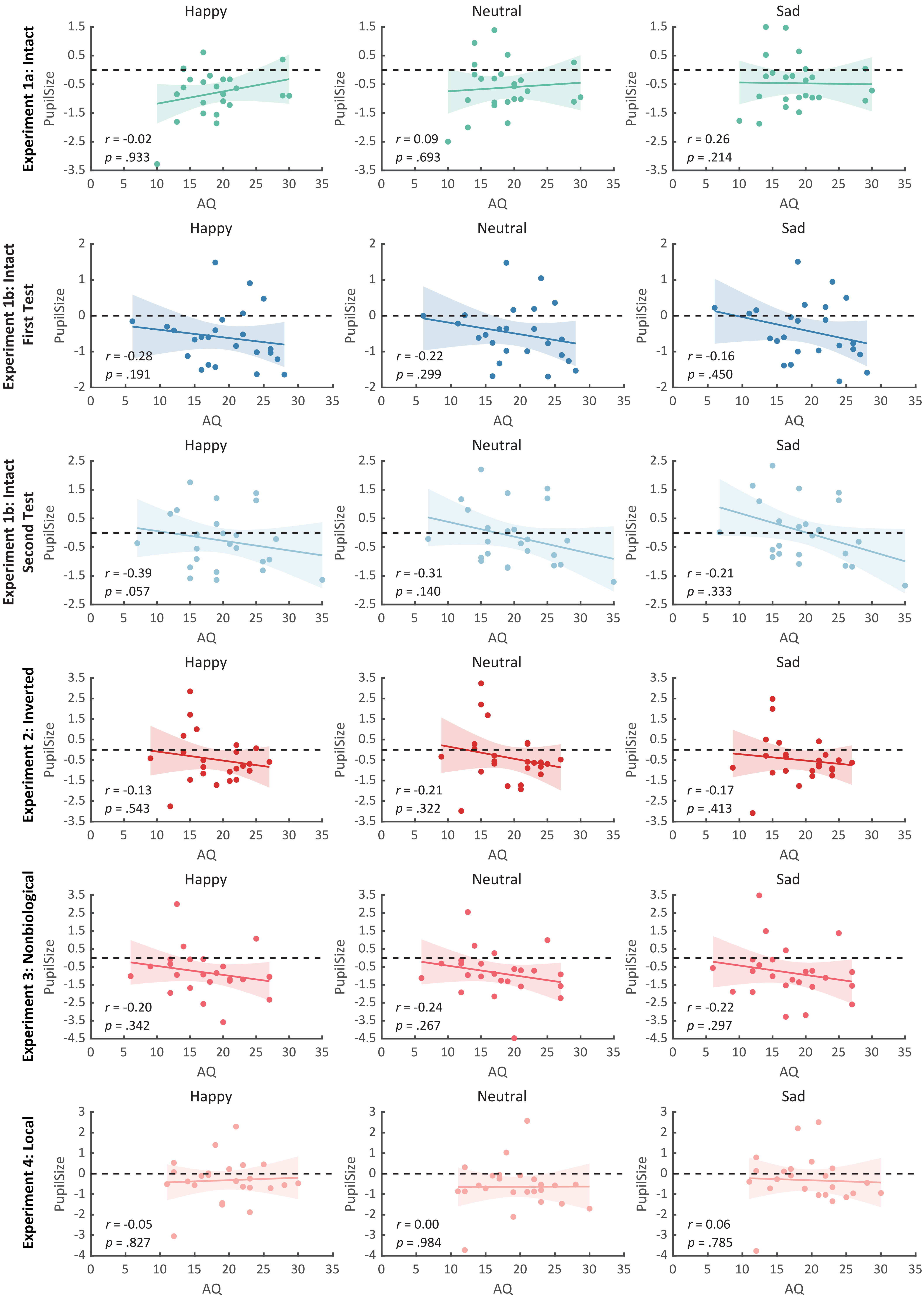
